# An Estradiol-Inducible Promoter Enables Fast, Graduated Control of Gene Expression in Fission Yeast

**DOI:** 10.1101/092338

**Authors:** Makoto J. Ohira, David G. Hendrickson, R. Scott McIsaac, Nicholas Rhind

## Abstract

The fission yeast *Schizosaccharomyces pombe* lacks a diverse toolkit of inducible promoters for experimental manipulation. Available inducible promoters suffer from slow induction kinetics, limited control of expression levels and/or a requirement for defined growth medium. In particular, no *S. pombe* inducible promoter systems exhibit a linear dose response, which would allow expression to be tuned to specific levels. We have adapted a fast, orthogonal promoter system with a large dynamic range and a linear dose response, based on β-estradiol-regulated function of the human estrogen receptor, for use in *S. pombe*. We show that this promoter system, termed Z_3_EV, turns on quickly, can reach a maximal induction of 20 fold, and exhibits a linear dose response over its entire induction range, with few off target effects. We demonstrate the utility of this system by regulating the mitotic inhibitor Wee1 to create a strain in which cell size is regulated by β-estradiol concentration. This promoter system will be of great utility for experimentally regulating gene expression in fission yeast.

## INTRODUCTION

Cell biological studies often take advantage of inducible promoters, ideally ones that are orthogonal to the system of interest. However, the inducible promoters available for the fission yeast *Schizosaccharomyces pombe* all have significant drawbacks ((Forsburg 1993) and Table S1). The inducible promoter most frequently used in *S. pombe* research is the endogenous *nmt1* promoter and its variants (Maundrell 1993). The *nmt1* gene has a strong promoter, which is regulated by thiamine, and has been engineered into a series of variants with low to high expression levels (Basi et al. 1993). The disadvantages of the *nmt1* promoter include a high constitutive activity, even in its weakest version, pleiotropic effects of induction by thiamine removal, and a long, 16-hour induction time. A synthetic *nmt1*-based promoter repressed by lacO improves the kinetic performance (Kjærulff & Nielsen 2015). Another synthetic promoter strategy combined tetO with the CaMV 35S promoter (Faryar & Gatz 1992; Erler et al. 2006). However, tetO/CaMV expression is comparable to only the weakest *nmt1* promoter variant and also takes between 9 to 12 hours to induce (Forsburg 1993; Erler et al. 2006). A more recent version of the tetO system has improved its expression kinetics (Zilio et al. 2012). Nonetheless, mitochondrial toxicity of tetracycline remains a concern (Moullan et al. 2015). The endogenous *urg1* promoter has been shown to have favorable induction and repression kinetics (Watt et al. 2008), but it suffers from the disadvantages of being regulated by uracil (which requires growth in defined media and has other transcriptional effects), working best only at its endogenous locus (Watson et al. 2013), and not having a well-characterized dose response.

McIsaac *et al.* recently designed and implemented a synthetic promoter system, named Z_3_EV, in *Saccharomyces cerevisiae*. This system is characterized by rapid and tunable induction of a desired target gene with the exogenous mammalian hormone β-estradiol. In *S. cerevisiae*, this system was shown to have a large dynamic range with minimal off-target activity (McIsaac et al. 2013; McIsaac et al. 2014). To increase the array of gene expression tools available in fission yeast, we have adapted the Z_3_EV system for use in *S. pombe*.

The Z_3_EV artificial transcription factor is a fusion of three domains: the Zif268 zinc-finger DNA binding domain containing three zinc fingers, the human estrogen receptor ligand-binding domain, and the VP16 viral activator domain (McIsaac et al. 2013; McIsaac et al. 2014). The estrogen receptor is an effective allosteric switch: binding of β-estradiol to the estrogen receptor disrupts its interaction with Hsp90 and causes rapid nuclear import of the Z_3_EV protein. The Z_3_EV-responsive promoter (Z_3_EVpr) consists of six Zif268 DNA binding sequences inserted into a modified *S. cerevisiae* GAL1 promoter from which the canonical Gal4 binding sites have been removed. Target genes placed immediately downstream of Z_3_EVpr in a strain expressing Z_3_EV become conditionally expressed upon β-estradiol addition to the culture. A GFP reporter driven by the Z_3_EV system in *S. cerevisiae* plateaued at about ~10-fold fluorescence over background when induced with 100 nM β-estradiol (McIsaac et al. 2013). GFP mRNA increased to between 50-100 fold over background approximately 20 minutes after induction with 1 μM of β**-**estradiol (McIsaac et al. 2013; McIsaac et al. 2014). Though *S. cerevisiae* and *S. pombe* diverged ~420 million years ago (Sipiczki 2000), the conservation of Hsp90 (which is present in all organisms except Archaea (Gupta 1995)) suggested the Z_3_EV system may also function in *S. pombe*.

In this manuscript, we describe the successful adaptation of the Z_3_EV system for use in *S. pombe* and a set of constructs to make its implementation straightforward. We find that the Z_3_EV system enables rapid, titratable induction of target gene expression in *S. pombe.* Increasing β-estradiol concentration results in a proportional increase of target gene expression. We use RNA-seq to show that the system has few off-target effects and thus can be used to regulate genes of interest with minor perturbations to cell physiology. Finally, we demonstrate that this system enables the study of native gene function: induction of the cell cycle regulator Wee1 by Z_3_EV results in dose-dependent increase in cell size. Collectively, these results highlight the utility of Z_3_EV to flexibly control gene expression in *S. pombe* without the significant drawbacks of previous synthetic systems.

## MATERIALS AND METHODS

### Media and growth conditions

We used standard fission yeast culture and growth conditions (YES media, 30°C) as previously described (Forsburg & Rhind 2006). β-estradiol (E2758, Sigma Aldrich) was dissolved in ethanol at 10mM and stored at −20°C. β-estradiol was added to media from this 10mM stock to the final concentration. An equivalent volume of ethanol was added to untreated control cultures.

### Strain and Plasmid construction

Strains and plasmids used in this study are listed in Tables S2 and S3. To create the Z_3_EV protein integration plasmid (pFS461), we used Gibson assembly (NEB) to join two fragments: the Z_3_EV CDS was amplified from DBY19011 (McIsaac et al. 2014) using primers MO198 and MO199 (Table S4), the *leu1* integrating vector (pFS181) (Sivakumar et al. 2004), which contains a strong constitutive *adh1* promoter, was amplified using primers MO197 and MO200. pFS461 was cleaved uniquely within the *leu1* ORF using XhoI and transformed into parent strain yFS110 to yield the transformant yFS949.

The plasmid containing GFP-NLS expressed from the Z_3_EV promoter (pFS462) was created by Gibson assembly (NEB) of four fragments: the Z_3_EV promoter sequence was amplified from pMN9 (McIsaac et al. 2013) using primers MO219 and MO220, the *his7* integration vector was amplified from pBSSK-His7(F) (Apolinario et al. 1993) using MO218 and MO225, the GFP-NLS CDS was amplified from pSO745 (courtesy of Snezhana Oliferenko) using MO221 and 222 and inserted up-stream of the Z_3_EV promoter; the *leu1* C-terminal sequence was amplified from *S. pombe* genomic DNA using MO223 and MO224, and appended downstream of the GFP-NLS sequence. pFS462 was cleaved uniquely within the *his7* ORF using AflII (NEB) and transformed into yFS949, to yield transformant strain yFS951. The *S. pombe* genome contains no potential Zif268 binding sites when searched with NCBI BLASTN under normal or nearest match specificity.

The plasmid containing beetle luciferase expressed from the Z_3_EV promoter (pFS465) was created by Gibson assembly (NEB) to replace GFP with luciferase in pFS462. The beetle luciferase ORF was amplified from pFS470 using primers MO236 and MO237; the pFS462 vector was amplified with MO235 and MO238. pFS465 was cleaved uniquely at the *his7* ORF using AflII and transformed into the Z_3_EV protein-expressing strain yFS948. This intermediate strain was crossed with a strain constitutively expressing renilla luciferase as a translational fusion with *ade4* (yFS871) to yield our dual-luciferase strain (yFS954).

### Flow cytometry

GFP fluorescence was quantified using a Becton Dickinson FACScan flow cytometer. Cells were harvested at specified time points after induction with β-estradiol, spun down at 1 kG for 5 minutes at 4°C, washed with water, quickly spun down at RT, and resuspended in PBS. For each sample, we recorded the green fluorescence signal in the FL1 channel from 10,000 cells and normalized the mean intensity to the 0 timepoint or 0 dose sample.

### RNA extraction and RT-qPCR

10 ODs of cells were harvested at specified time points after induction, spun down at 1 kG for 5 minutes at 4°C, washed with water, briefly spun down at RT, flash frozen in LN_2_, and stored at - 20°C. RNA was extracted from the cell pellet by bead beating with Trizol (Ambion) according to the manufacturer’s instructions, followed by isopropyl alcohol precipitation, ethanol wash, and stored at −20°C. The RNA was treated at a ratio of 1ug of RNA per unit of Amp-grade DNase I (Invitrogen) per the manufacturer’s instructions. Gene-specific cDNA was obtained using Superscript III (Invitrogen) per the manufacturer’s instructions and the following primers: MO233 for GFP, MO191 for *cdc2*, and LD8 for *ade4*. qPCR primers MO233 and MO221 yielded an amplicon length of 290nt within the GFP CDS. Primers MO192 and MO193 yielded an amplicon of 152nt within the cdc2 CDS. Primers LD9 and LD10 yielded amplicon size 163nt within the ade5 CDS. BLASTing these RT and qPCR primers against the *S. pombe* genome showed that they were either unique to the target gene, or in the case of GFP had no match within the genome and were unique to the GFP CDS. Specificity of the primers was shown via melt curve derivative, and standard curves were generated for all primer sets. qPCR was performed in 20 μL reactions containing 5μL of cDNA, 10μL 2X SYBR FAST qPCR Master Mix Universal (KK4601, KAPA Biosystems), 0.4μL of each 10μM primer and 4 μL water. Reactions were run in technical triplicates on a Bio-Rad CFX system, and Cq values were obtained via the Bio-Rad CFX Manager.

### Dual luciferase assay

Dual luciferase assay (Promega) was performed according to manufacturer’s instructions. Ten ODs of cells were suspended in 200 μL 1x Passive Lysis Buffer (PLB) and approximately 200 μL of zirconia/silica beads (Biospec). The cells were lysed by vortexing for 10 minutes at maximum RPM at 4°C. The lysate was spun at 16 kG at 4°C for 5 minutes. Triplicate 10 μL samples of cleared lysate were loaded into separate wells of a 96 well plate, each with 50 μL of pre-dispensed Luciferase Assay Reagent II. The luminometer was programmed for a 2 second rest and a 10 second measurement. 50μL of Stop and Glow Reagent was dispensed, followed by the same rest and measurement.

### Curve fitting

Dose-response data was fit with

Y = base+((max-base)*(x^slope/(x^slope+EC_50_^slope)))

Where base is the reported level at 0 dose, max is the reporter level at saturating dose, EC_50_ is the dose (effector concentration) at 50% reporter expression, and slope is the ratio of change in reporter expression relative to dose (DeLean et al. 1978).

Response-kinetics data was fit with

Y = base+((max-base)*(1-e^(-(t-lag)/half-life)))

Where base is the reporter level at t = 0, which is set to 1, max is the reporter level at the end of the tome course, lag is the time until the reported level begins to increase, and half-life is the half-life of the reporter. Note that this observed half-life combines both the degradation of the reporter and its dilution due to cell growth during the time course.

Fits were calculated in Igor Pro 6.37 (Wavemetrics).

### RNA-seq

10 ODs of cells were harvested before and after 3 hours of induction with 1 μM β-estradiol in three biologically independent replicates. RNA was extracted as described above. RNA quality was confirmed by denaturing gel and Bioanalyzer (Agilent) analysis.

Ribosomal RNA was depleted from 1 μg of total RNA using Ribo-Zero™ Yeast Magnetic Gold (Illumina), followed by a cleanup using Agencourt RNAClean XP beads (Beckman Coulter) and elution with 19.5 μL of Elute, Prime, Finish mix from the TruSeq stranded mRNA Sample Preparation Kit (Illumina). We performed library construction per the vendor’s instructions starting with cDNA synthesis. We pooled the resulting barcoded cDNA libraries and subjected them to 70 base pair paired-end sequencing on an Illumina NextSeq.

Sequencing reads were aligned to the Ensembl EF2 *S. pombe* genome assembly accessed from Illumina’s iGenome database <http://support.illumina.com/sequencing/sequencing_software/igenome.html> (modified to include GFP) and to the EF2 transcriptome (also modified to include GFP) using Tophat2 with the options I 10000 i 10 to constrain splice junction searching to < 10,000 bp and > 10 bp. We estimated transcript abundance and called differentially expressed genes using Cuffdiff2 with a false discovery rate of 5% (Trapnell et al. 2009). We filtered out genes with no counts or noisy signal (coefficient of variation > 0.5) in either the untreated and treated samples. Sequence data available at NIH GEO with accession number GSE92246.

Gene enrichment analysis was conducted on the AnGeLi server (Bitton et al. 2015).

## RESULTS

### Z_3_EV Functions in *S. pombe*

To adapt the Z_3_EV system to *S. pombe*, we expressed the Z_3_EV protein from the strong constitutive *adh1* promoter on a plasmid (pFS461) integrated at the *leu1* locus and expressed GFP-NLS from Z_3_EVpr on another plasmid (pFS462) integrated at the *his7* locus (Figure 1A). Upon addition of β-estradiol, Z_3_EV translocates into the nucleus and drives GFP expression (Figures 1B and 1C). This Z_3_EV-driven GFP system enabled us to confirm the function of the Z_3_EV system in *S. pombe* and quantitatively characterize its expression kinetics. Upon addition of 1 μM β-estradiol, we observed an approximately 4-fold increase in fluorescence signal over 5 hours of induction (Figure 1D). Fitting an induction-kinetics model to the data indicates that GPF takes about 1 hour to express, consistent with the slow maturation of GFP fluorescence (Heim et al. 1994). The fit also allows us to infer that the half-life of GFP in cells is about 2 hours. However, this observed half-life combines both the protein degradation rate and its dilution rate due to cell growth. The fact that the yeast doubling time of about 2 hours suffices to explain the observed GFP half-life suggests that the rate of GFP degradation is significantly slower that its rate of dilution. The range of GFP expression from Z_3_EVpr is within the range of endogenous gene expression and between the strengths of the wild type and medium strength *nmt1* promoters (Basi et al. 1993) (Figure 1E).

**Figure 1.**
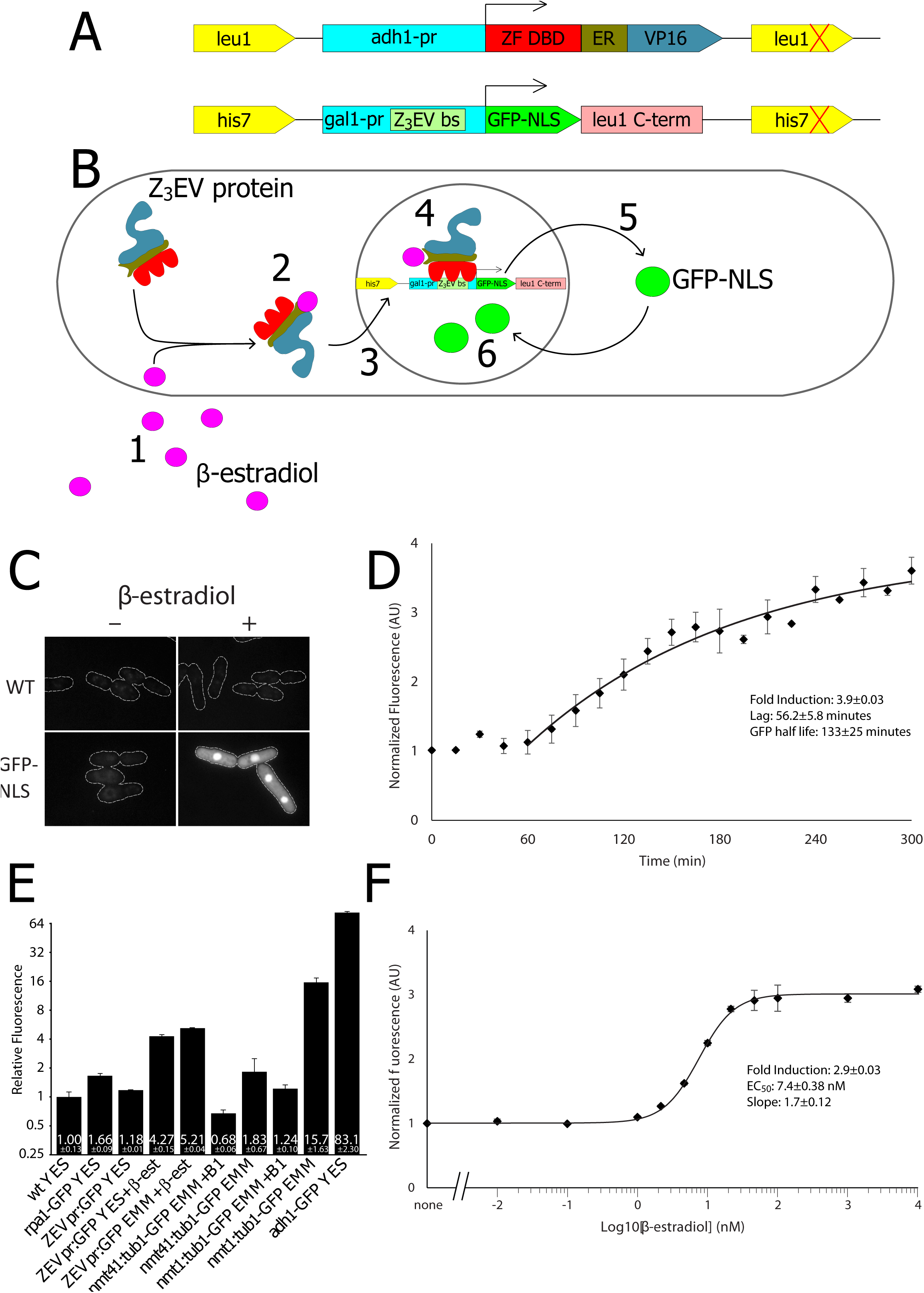
Z_3_EV Regulates Dose-Dependent Expression in *S. pombe*. (A) Schematic representation of the β-estradiol-induced promoter constructs. (B) Schematic representation of the β-estradiol-induced promoter system. The system was designed such that (1) added β-estradiol (2) binds to Z_3_EV protein leading to (3) translocation into the nucleus, (4) transcriptional activation of the GFP reporter gene, (5) GFP expression, and (6) translocation of GFP back into the nucleus. (C) Z_3_EV-dependent expression of GFP. Cells expressing Z_3_EV and containing the Z_3_EVpr:GFP expression cassette (yFS951) and wild-type cells (yFS110) were growth with or without 1 μM β-estradiol for 4 hours and imaged for green fluorescence. The outlines of the cells, taken from bright-field images, are also shown. (D) Z_3_EVpr:GFP induction kinetics. *adh1*:Z_3_EV Z_3_EVpr:GFP cells (yFS951) were induced with 1 μM β-estradiol and sampled every 20 minutes. Mean fluorescence of cells in the culture relative to uninduced cells was measured by flow cytometry. n=3; the mean and standard error bars are shown. The curve is an induction-kinetics model fit to the data from which the induction parameters were extracted. (E) Z_3_EVpr:GFP expression levels relative to other promoters. wt (yFS105), rpa1-GFP (yFS831), *adh1*:Z_3_EV ZEVpr:GFP (yFS951), nmt41:tub1-GFP (yFS386), nmt1:tub1-GFP (yFS387) and adh1-GFP (yFS313) cell were grown in the rich medium (YES) or synthetic medium (EMM). The ZEV promoter was induced with 1 μm β-estradiol. The *nmt1* promoter, or a moderate-strength mutant thereof (*nmt41*), was repressed with 15 μm thiamine (B1). Cells were assayed for mean fluorescence relative to wild-type background fluorescence by flow cytometry. n=3 for all strains; the mean and standard error bars are shown. (F) Dose response of Z_3_EVpr:GFP expression. *adh1*:Z_3_EV Z_3_EVpr:GFP cells (yFS951) were induced with various concentrations of β-estradiol, sampled after 5 hours and assayed for mean cellular fluorescence relative to uninduced cells by flow cytometry. n=3; the mean and standard error bars are shown. The curve is an sigmoidal dose-response model fit to the data from which the dose-response parameters were extracted.

### Z_3_EV Expression is Linearly Dependent on β-Estradiol

To characterize the dose response of Z_3_EV to β-estradiol in *S. pombe*, we assayed Z_3_EV-dependent expression over 4 logs of β-estradiol concentration. We observed only background fluorescence below 1 nM, and an upper plateau of approximately 4-fold induction at or above 100 nM (Figure 1F). The background fluorescence below 1 nM was similar in un-induced cells, cells expressing Z_3_EV protein without a reporter, and wild-type cells (Figures 1C and 1F). Between 1 nM and 100 nM β-estradiol, we observed a linear dose-response of GFP expression to β-estradiol concentration (Figure 1F). Fitting with a sigmoidal dose-response curve shows a low nanomolar EC_50_ (effector concentration at 50% response) and a slope (the ratio of change in response relative to dose) in the neighborhood of 1.

Although the GFP reporter allows rapid validation of the system and single-cell analysis of expression, the background auto-fluorescence of cells obscures the full dynamic range of Z_3_EV-dependent expression. To more accurately measure the dynamic range of the Z_3_EV system, we used a beetle luciferase reporter. Luciferase has excellent signal-to-noise characteristics and by using a Z_3_EV:beetle luciferase reporter strain constitutively expressing renilla luciferase (Voon et al. 2005), we have an internal control for dual-luciferase quantification experiments. Luciferase expression showed similar kinetic and dose-response profiles to the GFP reporter (Figures 2A,B). However, the dynamic range of the luciferase luminescence was much larger, with maximal expression over 20-fold greater than repressed levels (Figure 2B).

**Figure 2.**
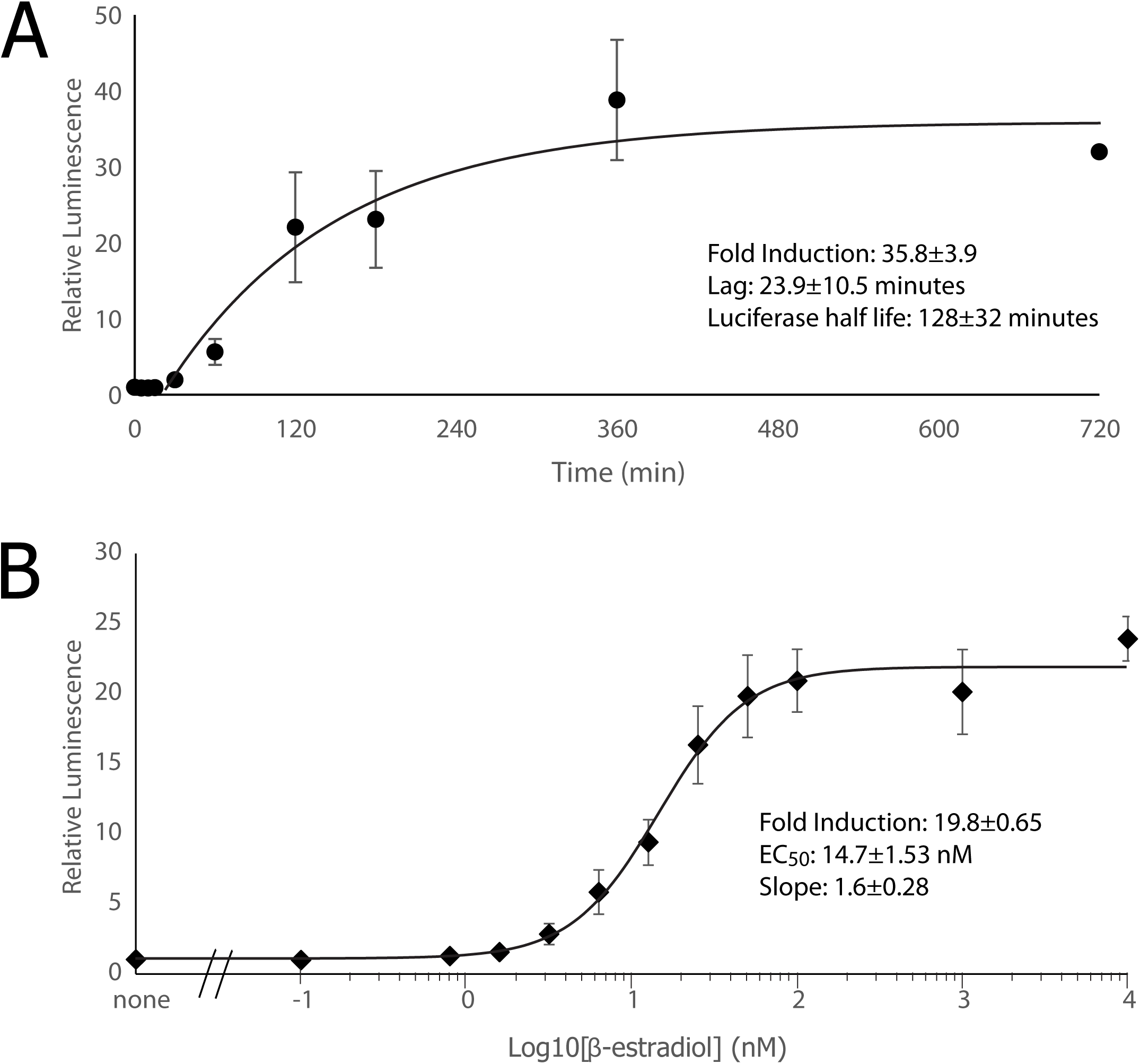
Z_3_EV Regulates Expression Over a Large Dynamic Range. (A) Z_3_EVpr:bLuc induction kinetics. Cells expressing Z_3_EV, containing the Z_3_EVpr:bLuc expression cassette and expressing a control Ade4-rLuc fusion (yFS954) were induced with 100 nM β-estradiol and sampled as indicated. Mean beetle luciferase activity of cells relative to renilla luciferase activity was measured by luminometry and normalized to uninduced cells. n=3; mean and standard error bars are shown. The curve is an induction-kinetics model fit to the data from which the induction parameters were extracted. (B) Dose response of Z_3_EVpr:bLuc expression. Z_3_EV Z_3_EVpr:bLuc Ade4-rLuc cells (yFS954) were induced with various concentrations of β-estradiol, sampled after 4 hours and assayed for mean beetle luciferase activity of cells relative to renilla luciferase activity by luminometry. Data was normalized to the uninduced sample. n=3 for all points except 720 minutes, for which n=1; mean and standard error bars are shown. The curve is an sigmoidal dose-response model fit to the data from which the dose-response parameters were extracted.

### mRNA Expression is Rapidly Induced by Z_3_EV Activation

To directly assay the transcriptional activity of Z_3_EVpr in response to Z_3_EV activation, we measured changes in reporter mRNA levels. A response-kinetics assay using RT-qPCR is similar to the luciferase assay with a 20-fold dynamic range peaking at 2 hours (Figure 3A). However, induction at the mRNA level is much faster than either of the reporter proteins. Induction of GFP transcripts was evident by 5 minutes after β-estradiol addition (Figure 3A). Fitting an induction-kinetics model to the data indicates that there in no measurable lag in mRNA expression and that the time taken to reach maximum expression is limited solely by the relatively long, ~40 minute, half-life of the GFP mRNA. Thus, the RT-qPCR-based assay provides a more sensitive measure of promoter activity than the fluorescence or luminescence-based assays, and demonstrates the rapid transcriptional response of the Z_3_EV system. Nonetheless, the dose-response parameters of all three assays are similar, with low nanomolar EC_50_s and slopes near 1.

**Figure 3.**
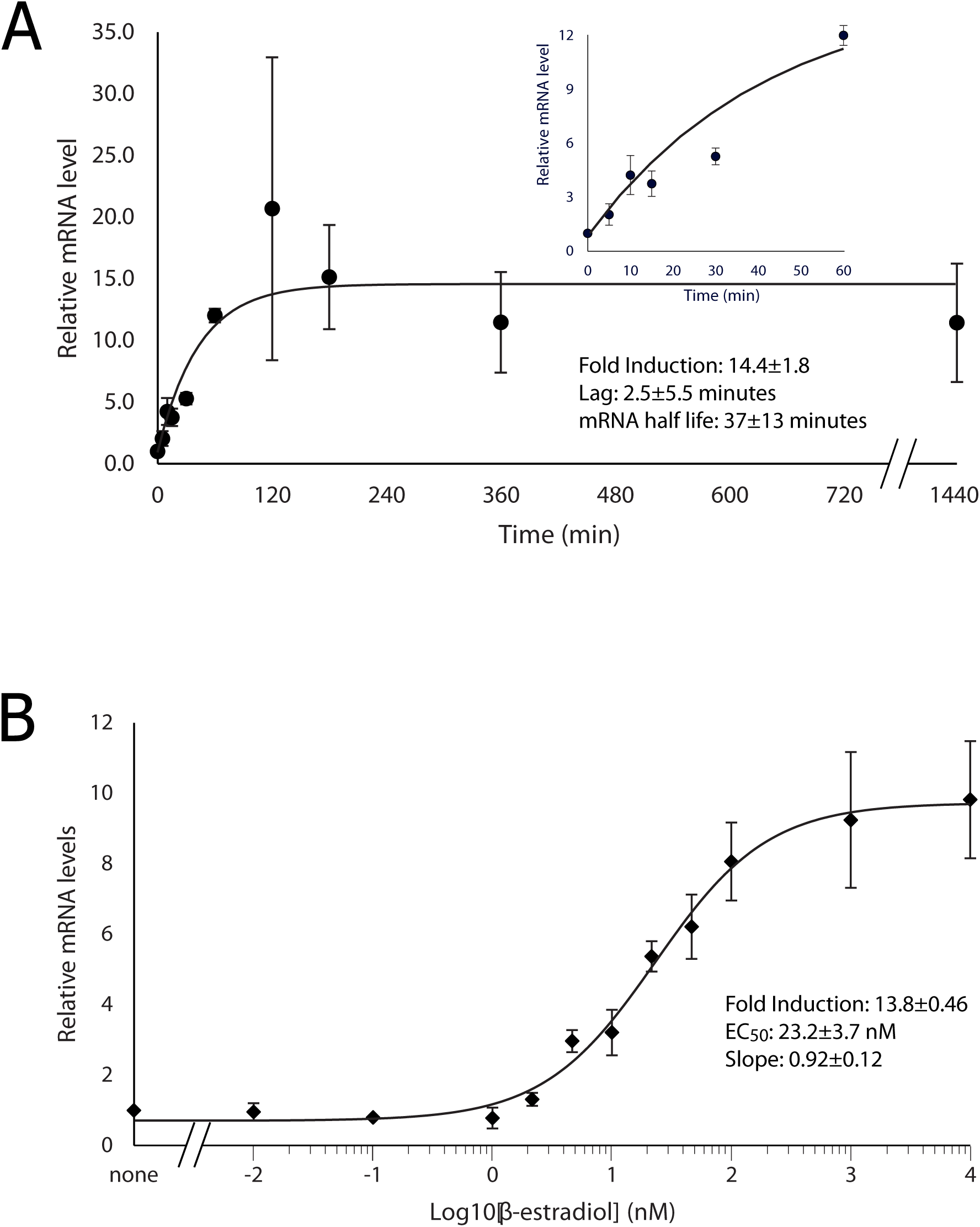
Z_3_EV Exhibits Rapid Induction Kinetics. (A) Z_3_EVpr:GFP mRNA induction kinetics. Z_3_EV Z_3_EVpr:GFP cells (yFS951) were induced with 1 μM β-estradiol and sampled as indicated. GFP mRNA levels relative to *cdc2* mRNA were measured by qPCR and normalized to uninduced cells. Early time points are shown in the inset graph. n=3; the mean and standard error bars are shown. The curve is an induction-kinetics model fit to the data from which the induction parameters were extracted. (B) Dose response of Z_3_EVpr:GFP mRNA expression. Z_3_EV Z_3_EVpr:GFP cells (yFS951) were induced with various concentrations of β-estradiol, sampled after 5 hours and assayed for GFP mRNA levels relative to *ade4* mRNA levels by qPCR. Data was normalized to the uninduced sample. n=3; the mean and standard error bars are shown. The curve is an sigmoidal dose-response model fit to the data from which the dose-response parameters were extracted.

### Growth effects of Z_3_EV Induction

There is some evidence that the VP16 domain can be toxic in yeast (Berger et al. 1992; Remacle et al. 1997; Silverman et al. 1994; McIsaac et al. 2011). To test the effect of the Z_3_EV protein (which includes a VP16 domain) on fitness, we compared the growth of wild type cells to cells expressing the Z_3_EV protein and to cells expressing both Z_3_EV protein and Z_3_EVpr-GFP under varying levels of β-estradiol induction.

Growth rates at various β-estradiol doses were measured and culture doubling times were calculated (Figure 4A). Wild-type cells are unaffected by addition of β-estradiol. In contrast, both the Z_3_EV protein-only expressing strain and the strain containing both Z_3_EV and a GFP reporter, showed a similar modest increase of doubling time with increasing β-estradiol (Figure 4A). However, the decrease in growth rate is less than 10% at 100 nM β-estradiol and even at 10 μM, where the reduction in growth rate is about 25%, the cells appear healthy by DIC microscopy. Since the EC_50_ for the reduction in growth rate is 10- to 30-fold higher than that for target-gene expression and the reduction only becomes substantial at concentrations above those required for maximal target-gene expression, the Z_3_EV system can be used over its full dynamic range with only a slight effect on growth rate.

**Figure 4.**
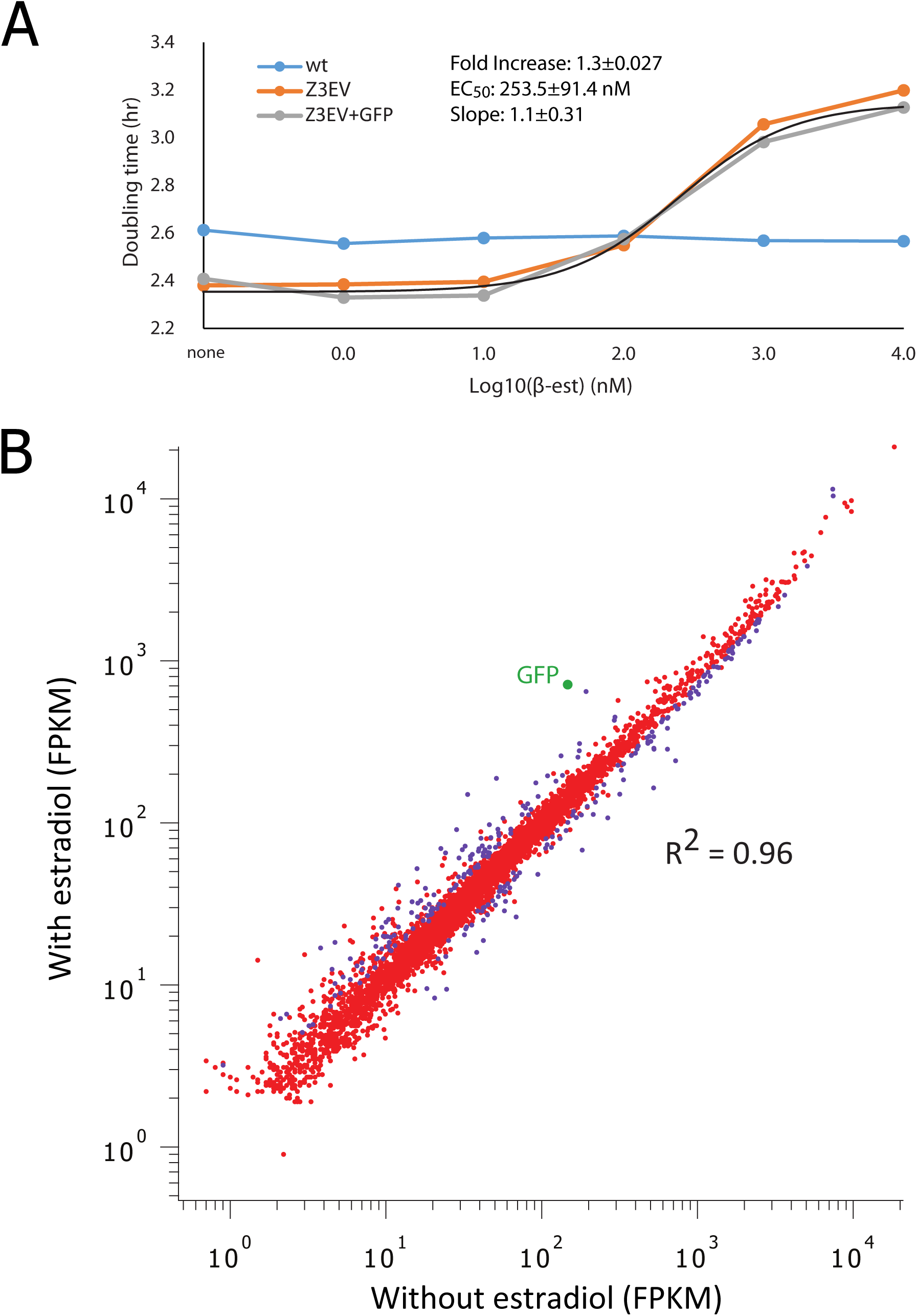
Global Effects of Z_3_EV Activity on Cell Growth and Gene Expression. (A) Activation of Z_3_EV causes a modest reduction in growth rate. Growth curves of wild-type (yFS110), Z_3_EV (yFS949) and Z_3_EV Z_3_EVpr:GFP (yFS951) strains at varying concentrations of β-estradiol. Growth was measured by optical density over 11 hours. The curve is an sigmoidal dose-response model fit to the data from which the dose-response parameters were extracted. The longer doubling time for the wild-type control at the lowest doses of β-estradiol may be due to the fact that our wild-type strain is auxotrophic for leucine, whereas both Z3Ez strains are leucine prototrophs. (B) Activation of Z_3_EV causes few off-target effects on mRNA expression. RNA-seq FPKM values of induced vs. un-induced Z_3_EV Z_3_EVpr:GFP cells (yFS951). GFP mRNA is highlighted in green.

### Z_3_EV has Few Off-Target Effects

To investigate the genome-wide effect of the β-estradiol induced Z_3_EV system in *S. pombe*, we used RNA-seq to profile the transcriptomes of our Z_3_EV strain under uninduced and induced conditions. Most genes remained unaffected by the induction of the Z_3_EV system with 1 μM of β-estradiol. Only 265 out of 5992 expressed loci showed greater than 1.32-fold change in expression after induction, a cutoff set so that genes reported as differentially expressed have a q value (FDR-corrected p value) of less than 0.05 (Table S5). Approximately equal numbers of genes showed an increase versus a decrease in expression. The largest increase was GFP mRNA, which showed a 4.9-fold increase (Figure 4B). The gene with the largest decrease in expression was *pho84*, at 3.2 fold.

Gene ontology enrichment analysis showed no significant enrichment within the set of genes that increased with Z_3_EV induction. The set of genes that decreased with Z_3_EV induction showed between 4-7 fold enrichment for various cytoplasmic translation or amino acid metabolic factors. It is unclear if the decrease in expression of genes associated with translation and amino acid biosynthesis is a cause or consequence of the reduced growth rate observed at 1 μM β-estradiol.

### Z_3_EV Regulation of *wee1* Expression Creates β-Estradiol-Dependent Cell Size Control

To demonstrate the experimental utility of the Z_3_EV promoter, we constructed a strain in which expression of the mitotic inhibitor Wee1 is under control of the Z_3_EV promoter. Wee1 is a dose-sensitive regulator of mitotic entry and thus cell size (Russell & Nurse 1987). We therefore expected that by regulating Wee1 expression, we would be able to regulate the size of cells at division. To do so, we replaced the endogenous *wee1* promoter with the Z_3_EV promoter in a strain expressing the Z_3_EV protein. In response to increasing doses of β-estradiol, cell size at division increases from 13 μm to 28 μm with a dose-response similar to those of the Z_3_EV-responsive fluorescent and luminescent reporters (Figure 5).

**Figure 5.**
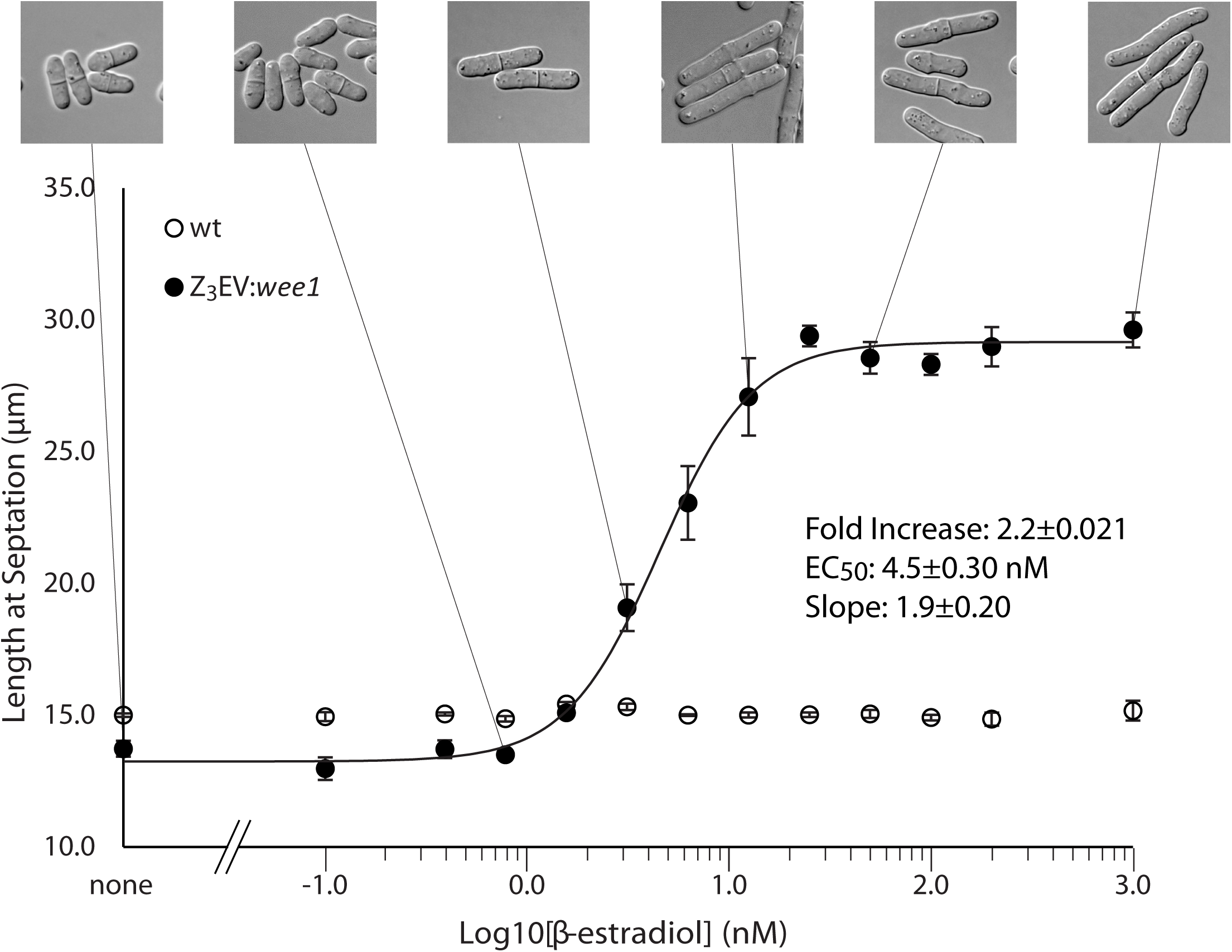
Z_3_EV-Dependent Regulation of Cell Size. (A) Z_3_EVpr:*wee1* regulates cell size in response to β-estradiol. Wild-type cells (yFS105) and cells expressing Z_3_EV and Z_3_EVpr:*wee1* (yFS970) were grown for 6 hours in varying concentrations of β-estradiol. (B) Quantitation of the length of cells at septation of cultures shown in 5A. The curve is an sigmoidal dose-response model fit to the data from which the dose-response parameters were extracted.

## DISCUSSION

Research using the fission yeast *S. pombe* has been hampered by the lack of convenient systems for regulation of gene expression. An ideal system would have a large dynamic range over which expression could be controlled by varying inducer concentration, would be rapidly induced in response to addition of inducer, would be based on an exogenous regulatory system, so as to have minimal incidental transcriptional effects, and would function in rich media, for cost efficiency. To establish such a system, we implemented a β-estradiol-regulated transcription system based on the Z_3_EV synthetic transcription factor, which was originally developed for use in *S. cerevisiae*, and showed that it operates successfully in *S. pombe*.

Maximal Z_3_EV activation by saturating levels of β-estradiol (> 100 nM) induces about a 20-fold increase in transcription from genes driven by Z_3_EVpr, the Z_3_EV responsive promoter (Figures 2A,B and 3A,B). This dynamic range compares favorably with other inducible promoters in *S. pombe*, which mostly show between 10- and 25-fold induction (Forsburg 1993; Watt et al. 2008), with the exception of the full-strength *nmt1* promoter which can induce up to 300-fold, often resulting in pathological overexpression.

The level of expression from Z_3_EVpr is within the range of normal *S. pombe* gene expression, with its uninduced level being just less than the *wee1* promoter and its maximal induced level being about 30% of maximal *nmt1* expression (Figures 1E and 5). However, the absolute level of expression from any promoter will depend on the stability of the target gene’s message and protein product. For applications that require lower basal expression levels, gene expression could be modified by adding mRNA or protein destabilizing elements (Voon et al. 2005), as has been done with the *urg1* promoter (Watson et al. 2013).

One of the major drawbacks of the *nmt1* promoter—the most commonly used promoter in *S. pombe* experiments—is its slow activation kinetics, requiring 16 hours to induce. Z_3_EV induces quickly, with mRNA expression evident at 5 minutes after induction (Figure 3A). Modeling of its induction kinetics suggests that there is no measurable lag in mRNA expression. The lag seen in the expression the GFP and luciferase reporter proteins are presumably due to their slow maturation kinetics. The modeling further suggests that the time to reach maximum mRNA expression is determined solely by the half-life of the target mRNA.

A major advantage of the Z_3_EV system over other expression systems available in *S. pombe* is that it has a linear dose response over the full range of its induction (Figures 1F, 2B, 3B and 5B). By titrating the concentration of β-estradiol, it is possible to obtain any expression level within its range, making the system convenient for controlling expression levels and allowing levels to be changed at will, affording tremendous experimental flexibility. Other *S. pombe* promoter systems have been modified to produce series of promoters of varying strengths (Basi et al. 1993; Watson et al. 2013). However, this approach allows access to only a fixed number of expression levels and does not allow flexible control of expression within one strain. The linear does response of Z_3_EV, in combination with the use of mRNA and protein destabilization elements to modify the absolute range of Z_3_EVpr-driven expression for any particular target gene, will afford a wide range of expression levels for any target protein (Voon et al. 2005; Watson et al. 2013).

Another significant advantage of the Z_3_EV system is that, being entirely foreign to *S. pombe*, it has very few off-target transcriptional effects. Only 265 genes show statistically significant chances in expression at the q < 0.05 level, which corresponds to a change of greater that 1.32 fold, and only 33 genes had a greater that 2-fold change in expression (Table S5). The statistically significantly up regulated genes did not show significant enrichment in any particular classes of genes. The down regulated genes showed enrichment for various cytoplasmic translation or amino acid metabolic factors. Since these genes are not enriched for even weak matches to the Z_3_EV binding site, it seems unlikely that they are direct targets. Instead their down regulation may reflect the slight reduction in growth rate at 1 μM β-estradiol (Figure 4A).

Even though activation causes only slight off-target changes in gene expression, it does have modest effects on cellular growth rates (Figure 4A). At 100 nM β-estradiol, the lowest concentration that induces maximal expression from Z_3_EVpr, cells double 10% slower than untreated cells; at 10 μM, they grow 25% slower, although they appear healthy. The presence of only minor off-target effects, along with the fact that growth rate continues to decline at doses above those required for maximal specific transcription induction, suggests that the mild growth defect induced by β-estradiol is not due to specific transcriptional effects but may be due to non-specific disruption of chromatin structure.

A final advantage of the Z_3_EV system is that it does not require specific media and works in both rich and synthetic medium (Figure 1E). This fact allows for increased experimental flexibility and is economically advantageous relative to promoters that require synthetic media.

The utility of the Z_3_EV system in *S. pombe* is demonstrated by its ability to regulate cell size when used to drive expression of the mitotic inhibitor Wee1. Wee1 is a protein kinase that negatively regulates Cdc2, the master regulator of mitosis (Nurse 1990). Wee1 is a dose dependent regulator of entry into mitosis and thus the level of Wee1 expression controls the size of cells at division. (Russell & Nurse 1987). We replaced the *wee1* promoter with Z_3_EVpr and showed that we could regulate cell size at division from 13 μm to 28 μm by titrating β-estradiol concentration (Figure 5). Although a number of strategies have been previously used to elongate cells, these approaches use disruptive treatments, such as temperature shift or checkpoint activation, and generally result in acute G2 arrest, so arbitrary intermediate cell sizes have not been experimentally accessible. Thus, beyond being a simple demonstration of the utility of the Z_3_EV system in *S. pombe*, this strain will be a useful tool for the study for cell size biology.

Given the limited existing repertoire of promoters available for the *S. pombe*, and the multiple advantages of the Z_3_EV system, we believe that it will prove to be an invaluable tool for future functional studies in this important model organism.

## AUTHOR CONTRIBUTIONS

Conceptualization: MO,NR; Methodology: MO,DGH,RSM,NR; Formal Analysis: MO,DGH,RSM,NR; Investigation: MO,DGH; Data Curation: RSM,NR; Writing – Original Draft: MO,NR; Writing – Review & Editing: MO,DGH,RSM,NR; Visualization: MO,DGH,RSM,NR; Supervision: RSM,NR; Project Administration: NR; Funding Acquisition: RSM,NR

## ACKNOWLEDGEMENTS

We thank Snezhana Oliferenko for the GFP-NLS construct, Dan Keifenheim for the luciferase strains, members of the Rhind lab for helpful suggestions throughout this project, Tony Carruthers for help with kinetic curve fitting, and Damien Coudreuse and Sarah Dixon for helpful comments on the manuscript.

## SUPPLEMENTAL INFORMATION

**Table S1:** S. pombe promoter systems

| Promoter | Characteristics | Reference |
| --- | --- | --- |
| adh1 | Strong, constitutive | (Russell & Hall 1983; Russell & Nurse 1984) |
| ef1a-c | Strong, constitutive | (Matsuyama et al. 2008) |
| hCGα | Strong, constitutive | (Toyama & Okayama 1990) |
| hCMV | Strong, constitutive | (Toyama & Okayama 1990) |
| hAd E2 early | Strong, constitutive | (Swaminathan et al. 1993) |
| HIV-1 LTR | Strong, constitutive | (Toyama et al. 1992) |
| hHSP70 | Strong, constitutive | (Prentice & Kingston 1992) |
| isd90 | Strong, constitutive | (Verma et al. 2014) |
| tif51 | Strong, constitutive | (Matsuyama et al. 2008) |
| cam1 | Weak, constitutive | (Matsuyama et al. 2008) |
| SV40 | Weak, constitutive | (Forsburg 1993) |
| pho1 | Adenine repressible | (Schweingruber et al. 1992) |
| fbp1 | Glucose repressible | (Hoffman & Winston 1989) |
| inv1 | Glucose repressible | (Iacovoni et al. 1999) |
| ctr4 | Copper repressible | (Bellemare et al. 2001) |
| hsp16 | Stress inducible | (Fujita et al. 2006) |
| tetO/CaMV | Weak, tet inducible | (Forsburg 1993; Faryar & Gatz 1992; Erler et al. 2006) |
| tetO<sub>7</sub>-TATA<sub>CYC1</sub> | Medium tet inducible | (Zilio et al. 2012) |
| nmt1 | Weak-Strong inducible | (Basi et al. 1993) |
| nmt1-ts | Truncated, ts nmt1 | (Kumar & Singh 2006) |
| nmt1-LacO | LacI repressed nmt1 | (Kjærulff & Nielsen 2015) |
| urg1 | Medium uracil inducible | (Watt et al. 2008) |

**Table S2:** Strains

| Strain | Genotype | Reference |
| --- | --- | --- |
| yFS109 | h+ leu1-32 ura4-D18 ade6-210 his7-366 | lab stock |
| yFS110 | h- leu1-32 ura4-D18 ade6-210 his7-366 | lab stock |
| yFS313 | h- leu1-32 ura4-D18 ade6-210 his7-366 adh1:GFP(KanMX) | lab stock |
| yFS386 | h- leu1-32 ura4-D18 nmt41-tubA-GFP | lab stock |
| yFS387 | h- leu1-32 ura4-D18 nmt1-tubA-GFP | lab stock |
| yFS831 | h- leu1-32 ura4-D18 rpa1-GFP-hphMX6 | lab stock |
| yFS871 | h- leu1-32 ura-D18 ade4-rLuc(NatMX) | lab stock |
| yFS945 | h- ura4-D18 leu1::pFS461 | this paper |
| yFS948 | h+ ura4-D18 ade6-210 his7-366 leu1::pFS461 | this paper<sup>1</sup> |
| yFS949 | h- ura4-D18 ade6-210 his7-366 leu1::pFS461 | this paper<sup>1</sup> |
| yFS951 | h- ura4-D18 ade6-210 leu1::pFS461 his7::pFS462 | this paper<sup>1</sup> |
| yFS954 | h+ ura4-D18 ade6-210 leu1::pFS461 his7::pFS465 ade4-rLuc(NatMX) | this paper |
| yFS970 | h- ura4-D18 leu1-32::pFS461 Z<sub>3</sub>EVpr:wee1(KanMX) | this paper<sup>1</sup> |

1) These strains will be available from NRBP http://yeast.lab.nig.ac.jp/nig/index_en.html.

**Table S3:**
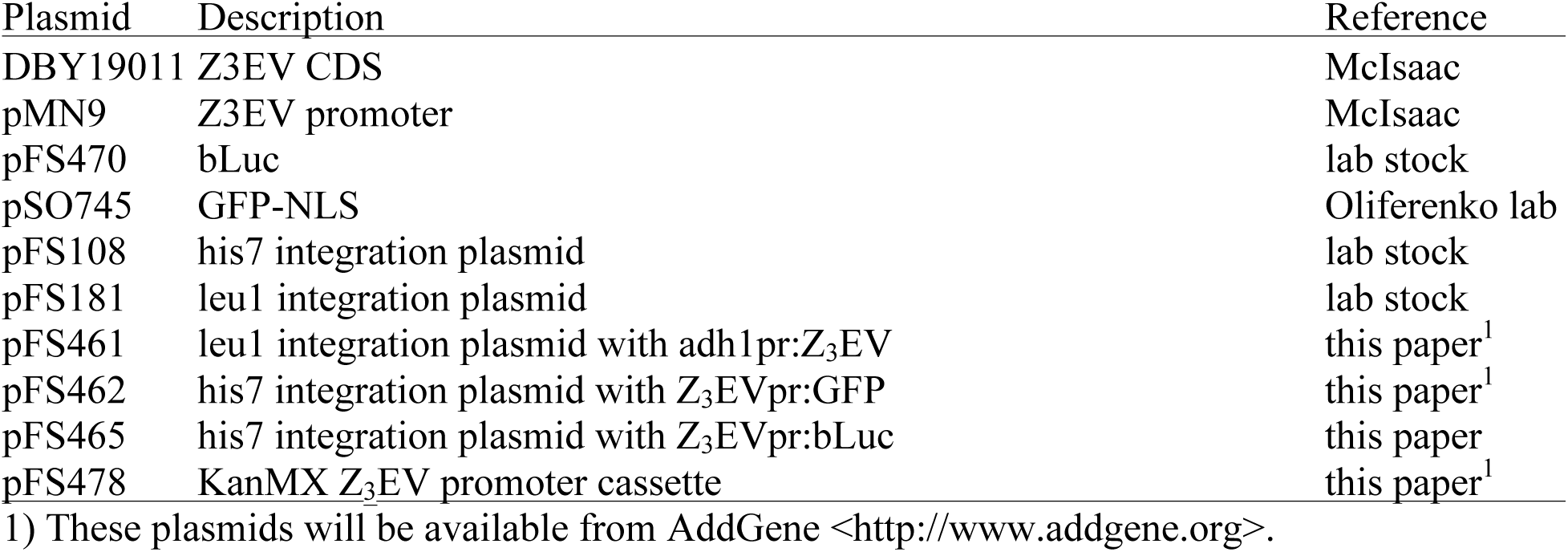
Plasmids

| Plasmid | Description | Reference |
| --- | --- | --- |
| DBY19011 | Z3EV CDS | McIsaac |
| pMN9 | Z3EV promoter | McIsaac |
| pFS470 | bLuc | lab stock |
| pSO745 | GFP-NLS | Oliferenko lab |
| pFS108 | his7 integration plasmid | lab stock |
| pFS181 | leu1 integration plasmid | lab stock |
| pFS461 | leu1 integration plasmid with adh1pr:Z3EV | this paper<sup>1</sup> |
| pFS462 | his7 integration plasmid with Z3EVpr:GFP | this paper<sup>1</sup> |
| pFS465 | his7 integration plasmid with Z3EVpr:bLuc | this paper |
| pFS478 | KanMX Z3EV promoter cassette | this paper<sup>1</sup> |

1) These plasmids will be available from AddGene <<http://www.addgene.org>>.

**Table S4:**
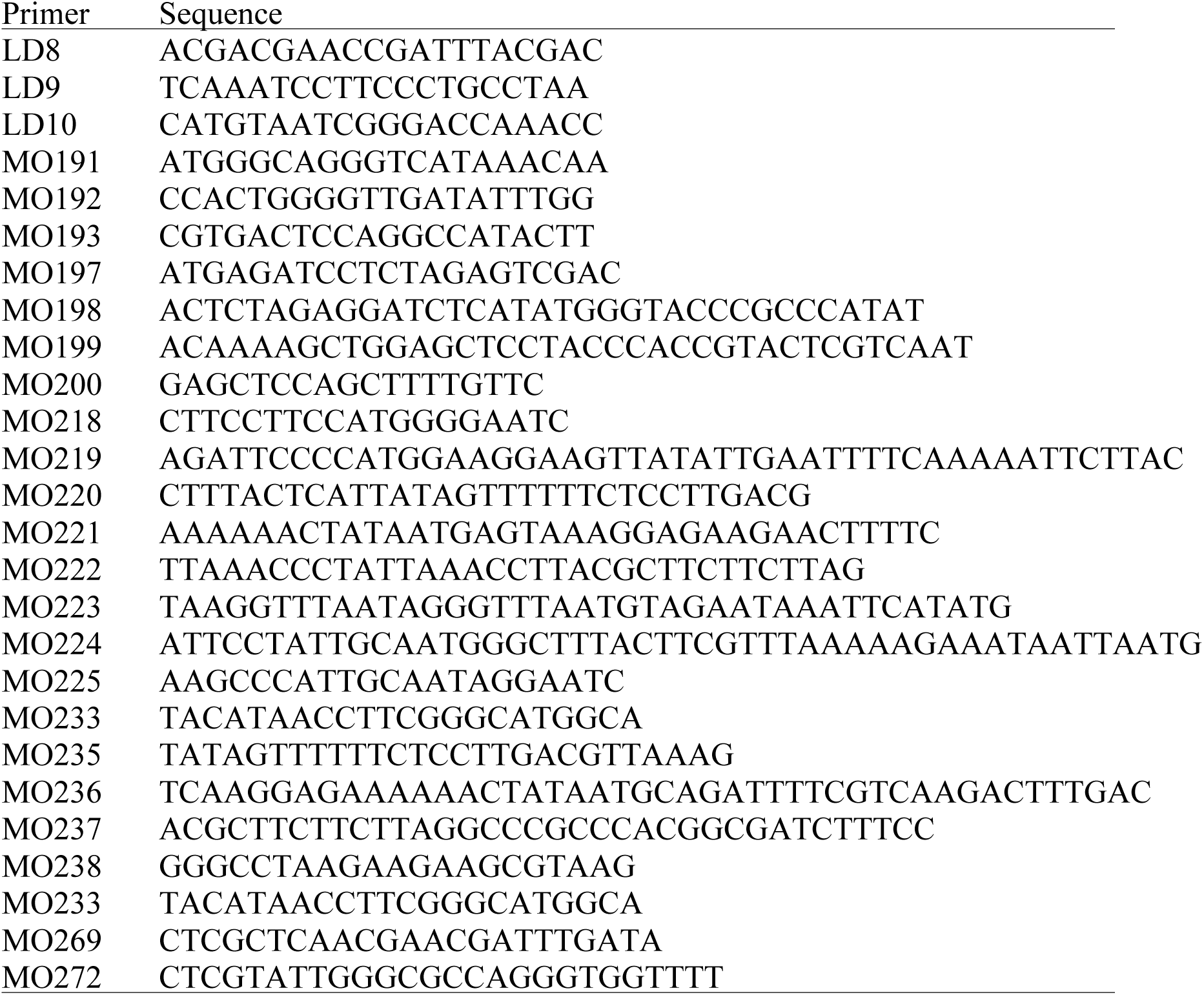
Primers

**Table S5:** MRNA expression changes upon Z_3_EV activation The accompanying workbook contains differential expression analysis of the RNA-seq data presented in Figure 4B. The first worksheet contains the average normalized expression level for each gene in the *S. pombe* genome, along with statistical analyses. The second spreadsheet contains the normalized data for each of the three biological replicates.

## REFERENCES

Apolinario, E. et al., 1993. Cloning and manipulation of the Schizosaccharomyces pombe his7 +gene as a new selectable marker for molecular genetic studies. Current Genetics, 24(6), pp.491–495.

Basi, G., Schmid, E. & Maundrell, K., 1993. TATA box mutations in the Schizosaccharomyces pombe nmtl promoter affect transcription efficiency but not the transcription start point or thiamine repressibility. Elsevier Science Publishing, 123(1), pp.131–136.

Bellemare, D.R. et al., 2001. A novel copper-regulated promoter system for expression of heterologous proteins in Schizosaccharomyces pombe. Gene, 273(2), pp.191–198.

Berger, S.L. et al., 1992. Genetic isolation of ADA2: a potential transcriptional adaptor required for function of certain acidic activation domains. Cell, 70(2), pp.251–265.

Bitton, D.A. et al., 2015. AnGeLi: A tool for the analysis of gene lists from fission yeast. Frontiers in Genetics, 6(NOV), pp.1–9.

DeLean, A., Munson, P.J. & Rodbard, D., 1978. Simultaneous analysis of families of sigmoidal curves: application to bioassay, radioligand assay, and physiological dose-response curves. The American journal of physiology, 235(2), pp.E97–E102.

Erler, A. et al., 2006. Recombineering reagents for improved inducible expression and selection marker re-use in Schizosacchromyces pombe. Yeast, 23, pp.813–823.

Faryar, K. & Gatz, C., 1992. Construction of a Tetracycline-inducible promoter in Schizosaccharomyces pombe. Current Genetics, 21, pp.345–349.

Forsburg, S.L., 1993. Comparison of Schizosaccharomyces pombe systems expression. Nucleic Acids Research, 21(12), pp.2955–2956.

Forsburg, S.L. & Rhind, N., 2006. Basic methods for fission yeast. Yeast, 23(3), pp.173–183.

Fujita, Y. et al., 2006. Heat shock-inducible expression vectors for use in Schizosaccharomyces pombe. FEMS Yeast Research, 6(6), pp.883–887.

Gupta, R.S., 1995. Phylogenetic analysis of the 90 kD heat shock family of protein sequences and an examination of the relationship among animals, plants, and fungi species. Mol Biol Evol, 12(6), pp.1063–1073.

Heim, R., Prasher, D.C. & Tsien, R.Y., 1994. Wavelength mutations and posttranslational autoxidation of green fluorescent protein. Proceedings of the National Academy of Sciences, 91(26), pp.12501–12504.

Hoffman, C.S. & Winston, F., 1989. A transcriptionally regulated expression vector for the fission yeast Schizosaccharomyces pombe. 413 Gene, 84, pp.473–479.

Iacovoni, J.S., Russell, P. & Gaits, F., 1999. A new inducible protein expression system in fission yeast based on the glucose-repressed inv1 promoter. Gene, 232(1), pp.53–58.

Kjærulff, S. & Nielsen, O., 2015. An IPTG-inducible derivative of the fission yeast nmt promoter. Yeast, 32, pp.469–478.

Kumar, R. & Singh, J., 2006. A truncated derivative of nmt1 promoter exhibits temperature-dependent induction of gene expression in Schizosaccharomyces pombe. Yeast, 23, pp.55–65.

Matsuyama, A., Shirai, A. & Yoshida, M., 2008. A series of promoters for constitutive expression of heterologous genes in fission yeast. Yeast, 25(5), pp.371–376.

Maundrell, K., 1993. Thiamine-repressible expression vectors pREP and pRIP for fission yeast. Gene, 123(1), pp.127–130.

McIsaac, R.S. et al., 2011. Fast-acting and nearly gratuitous induction of gene expression and protein depletion in Saccharomyces cerevisiae. Molecular biology of the cell, 22(22), pp.4447–59.

McIsaac, R.S. et al., 2014. Synthetic biology tools for programming gene expression without nutritional perturbations in Saccharomyces cerevisiae. Nucleic Acids Research, 42(6), pp.1–8.

McIsaac, R.S. et al., 2013. Synthetic gene expression perturbation systems with rapid, tunable, single-gene specificity in yeast. Nucleic Acids Research, 41(4), pp.1–10.

Moullan, N. et al., 2015. Tetracyclines disturb mitochondrial function across eukaryotic models: A call for caution in biomedical research. Cell Reports, 10, pp.1681–1691.

Nurse, P., 1990. Universal Control Regulating the Onset of M-phase.pdf. Nature, 344(6266), pp.503–508.

Prentice, H.L. & Kingston, R.E., 1992. Mammalian promoter element function in the fission yeast Schizosaccharomyces pombe. Nucleic Acids Res, 20(13), pp.3383–3390.

Remacle, J.E. et al., 1997. Three classes of mammalian transcription activation domain stimulate transcription in Schizosaccharomyces pombe. EMBO Journal, 16(18), pp.5722–5729.

Russell, P. & Nurse, P., 1984. c & 25 + Functions as an Inducer in the M itotic Control of Fission Yeast. Cell, 45, pp.145–153.

Russell, P. & Nurse, P., 1987. Negative regulation of mitosis by wee1+, a gene encoding a protein kinase homologue. Cell, 49, pp.559–567.

Russell, P.R. & Hall, B.D., 1983. The primary structure of the alcohol dehydrogenase gene from the fission yeast Schizosaccharomyces pombe. J Biol.Chem., 258(1), pp.143–149.

Schweingruber, M.E. et al., 1992. Regulation of pho1-encoded acid phosphatase of Schizosaccharomyces pombe by adenine and phosphate. Current genetics, 22(4), pp.289–92.

Silverman, N., Agapite, J. & Guarente, L., 1994. Yeast ADA2 protein binds to the VP16 protein activation domain and activates transcription. Proceedings of the National Academy of Sciences of the United States of America, 91(24), pp.11665–8.

Sipiczki, M., 2000. Where does fission yeast sit on the tree of life? Genome biology, 1(2), p.1011.1–1011.4.

Sivakumar, S. et al., 2004. In vivo labeling of fission yeast DNA with thymidine and thymidine analogs. Methods, 33(3), pp.213–219.

Swaminathan, S. et al., 1993. Activation of a dual adenovirus promoter containing nonconsensus TATA motifs in Schizosaccharomyces pombe: role of TATA sequences in the efficiency of transcription. Nucleic Acids Research, 21(11), pp.2737–2746.

Toyama, R., Bende, S.M. & Dhar, R., 1992. Transcriptional activity of the human immunodeficiency virus-1 LTR promoter in fission yeast Schizosaccharomyces pombe. Nucleic Acids Res, 20(10), pp.2591–2596.

Toyama, R. & Okayama, H., 1990. Human chorionic gonadotropin alpha and human cytomegalovirus promoters are extremely active in the fission yeast Schizosaccharomyces pombe. FEBS letters, 268(1), pp.217–221.

Trapnell, C., Pachter, L. & Salzberg, S.L., 2009. TopHat: Discovering splice junctions with RNA-Seq. Bioinformatics, 25(9), pp.1105–1111.

Verma, H.K. et al., 2014. High level constitutive expression of luciferase reporter by lsd90 promoter in fission yeast. PLoS ONE, 9(7), pp.1–10.

Voon, D.C. et al., 2005. Use of mRNA and protein-destabilizing elements to develop a highly responsive reporter system. Nucleic Acids Research, 33(3), pp.1–10.

Watson, A.T. et al., 2013. Optimisation of the Schizosaccharomyces pombe urg1 expression system. PLoS ONE, 8(12), pp.1–16.

Watt, S. et al., 2008. urg1: A uracil-regulatable promoter system for fission yeast with short induction and repression times. PLoS ONE, 3(1), pp.1–8.

Zilio, N., Wehrkamp-Richter, S. & Boddy, M.N., 2012. A new versatile system for rapid control of gene expression in the fission yeast Schizosaccharomyces pombe. Yeast, 29, pp.425–434.

